# Volatile compounds released by undamaged plants influence the adaptive growth strategies of neighboring plants

**DOI:** 10.1101/2025.08.15.670058

**Authors:** André Åbonde, Merlin Rensing, Jannicke Gallinger, Vasti Thamara Juárez-González, Iris Dahlin, Dimitrije Markovic, German Martinez, Velemir Ninkovic

## Abstract

Plants constantly emit volatile organic compounds (VOCs) that can influence the physiology and behavior of neighboring plants. While the role of stress-induced VOCs in mediating plant-plant interactions is well established, the ecological significance of constitutive VOCs from undamaged plants remains less understood. We demonstrate that barley plants can detect the growth rate of their undamaged neighbors through constitutive VOCs and respond by adjusting their trade-off between growth and induced defense. Exposure to volatiles from cultivars with slower or faster growth triggered distinct shifts in biomass accumulation and gene expression in receiver plants, whereas exposure to VOCs from cultivars with similar growth rates elicited negligible responses. These divergent patterns reflect a trade-off between growth and induced defense, consistent with adaptive responses to anticipated competition. Our findings reveal a previously unrecognized adaptive function of constitutive VOCs in mediating receiver growth based on neighbor growth rate, emphasizing their role in shaping plant-plant interactions.

## Introduction

Throughout their life cycle, plants continuously perceive and respond to environmental signals, adjusting their growth and development to prevailing conditions (Taiz *et al*., 2023). Among these signals, volatile organic compounds (VOCs) enable plants to interact with other organisms, including neighboring plants (Bouwmeester *et al*., 2019). VOCs provide a chemical ‘fingerprint’ of a plant’s physiological status, allowing neighbors to assess the emitter’s growth and biotic or abiotic stress (Effah *et al*., 2019; Ninkovic *et al*., 2021). While most research has focused on induced VOCs, released in response to stressors such as herbivore or pathogen attack and how they trigger defense responses in neighboring plants (Das *et al*., 2013; Bouwmeester *et al*., 2019; Brosset & Blande, 2022; Murali-Baskaran *et al*., 2022), far less is known about the functions of constitutive VOCs emitted by undamaged plants. Emerging evidence indicates that these constitutive VOCs can induce resistance in neighboring plants, making them less attractive to herbivores (Kheam *et al*., 2023; Markovic *et al*., 2025) and more appealing to natural enemies (Ninkovic & Pettersson, 2003; Ninkovic *et al*., 2011, 2016). Nonetheless, their functions beyond defense elicitation remain largely unexplored (Brosset & Blande, 2022).

Stressed-induced VOCs not only mediate interactions with other organisms but also influence the regulation of physiological processes within the plant, including defense activation and resource allocation (Brosset & Blande, 2022). Plant defense mechanisms include constitutive defenses, which are active at a basal level, and induced responses that are activated upon threat detection. Induced plant defenses involve the production of secondary metabolites and transcriptional reprogramming. These responses can be triggered by external signals, such as damage to protective tissues or VOCs from neighboring plants, as well as by internal signals mediated by hormonal pathways (Taiz *et al*., 2023).

Because the production of defensive metabolites diverts energy from primary metabolism, it often does so at the expense of growth and reproduction (Taiz *et al*., 2023). Plants therefore balance resource allocation between defense and growth, a fundamental aspect of plant ecology and physiology (Huot *et al*., 2014; Züst & Agrawal, 2017). Activation of defense pathways and their associated transcriptional reprogramming often result in reduced growth, a trend more commonly observed in stressful environments, whereas plants may prioritize growth and development over defense in favorable conditions (He *et al*., 2022). Understanding the transcriptional activation of the defense and growth pathways is key to inferring their effects at the physiological level. As such, genome-wide analysis with high-throughput sequencing provides insight into these allocation decisions by identifying genes activated under specific conditions (Wang *et al*., 2009). Despite this, the transcriptional changes induced by VOCs are not understood in detail.

Here, we test the hypothesis that plants can perceive and interpret the growth strategies of undamaged neighbors through their constitutive VOC profiles. We studied interactions among three barley (*Hordeum vulgare*) cultivars with contrasting growth rates (slow, intermediate and fast) and assessed their effects on biomass production and genome-wide gene expression in receiver plants. We show that the cultivars emit distinct VOC profiles that influence the growth-defense trade-off in their neighbors (Fig. 1). This reveals a novel adaptive function of constitutive VOCs, encoding information about a neighbor’s growth strategy and shaping resource allocation decisions in the receiver. These findings expand the ecological significance of VOCs beyond defense, positioning them as integral components of a dynamic information network through which plants monitor and respond to their local environment.

**Fig. 1.**
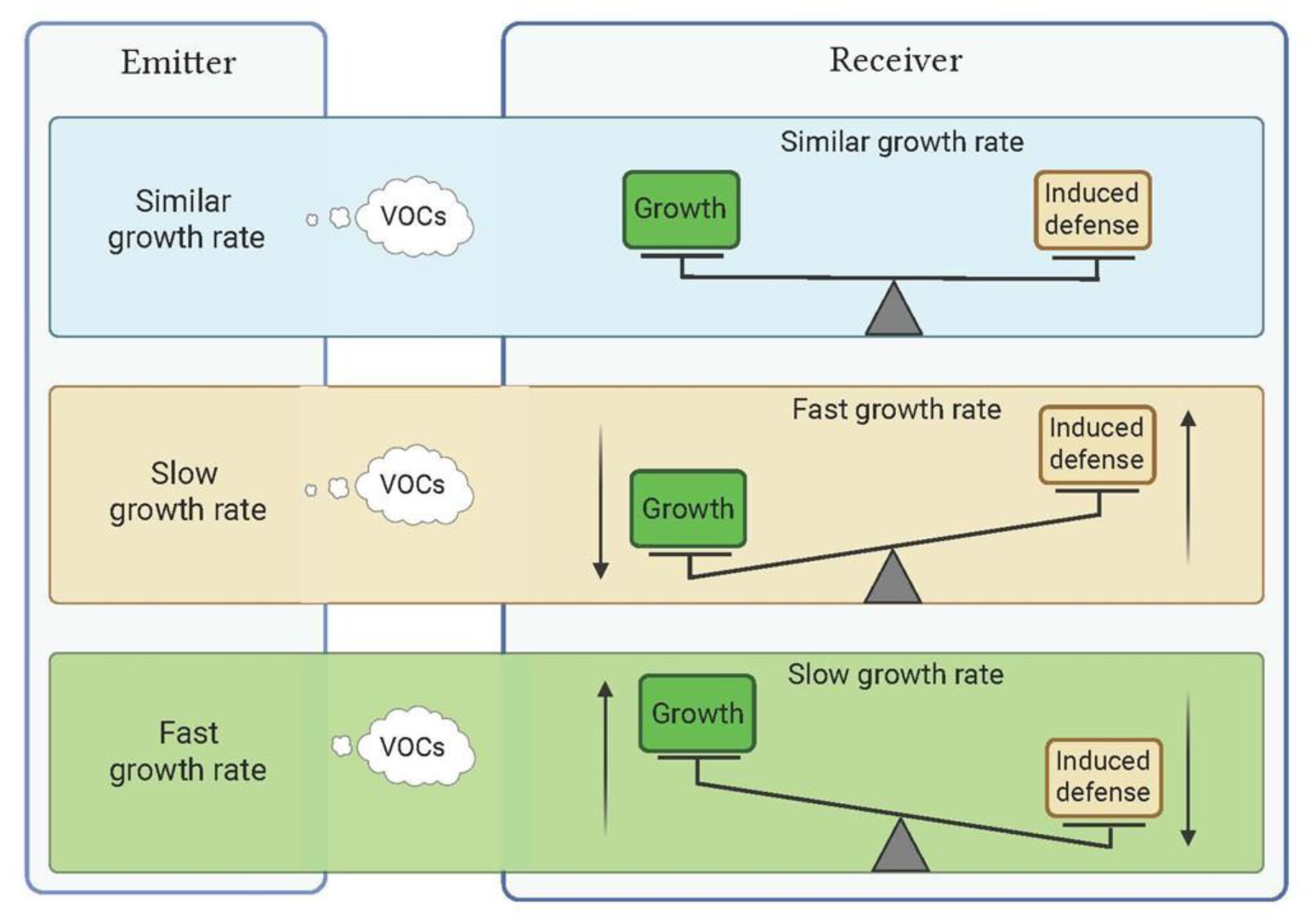
Constitutive VOCs emitted by undamaged emitter plants mediate adaptive growth strategies in receiver plants. Receivers with similar growth rates to the emitter remain unaffected. Fast-growing receivers reduce growth and allocate more to induced defense when exposed to VOCs from slow-growing emitters. In contrast, slow-growing receivers increase growth at the expense of induced defenses when exposed to VOCs from fast-growing emitters. (Created with BioRender.com).

## Materials and Methods

### Plant materials

Spring barley (*Hordeum vulgare* L.) cultivars Fairytale, Salome, and Luhkas (Scandinavian Seeds AB, Sweden) were selected based on their different growth strategies, previously shown to vary in growth rates (slow, intermediate, and fast, respectively) and to have genetically distinct pedigrees developed by breeders from different countries (Dahlin *et al*., 2020).

### Growing conditions

The climate in the growth chamber was maintained at 20±2°C, with a light regime of 16 hours of light and 8 hours of darkness (L16:D8), and 60% relative humidity. Light was provided by Light4Food lamps, situated 1 m above the table, with a central light strip (Philips LED, 77W, Poland) and side lights located to its left and right (Philips LED lighting IBRS, 55–60W, Netherlands), each following their own light regimen. Plants was grown in perforated plastic tubes (90 × 5 cm) filled with clean sand (Silversand 55, Sibelco Europe, Mölndal, Sweden). These tubes were positioned beneath the bench, with the open end of each tube passed through a 5 x 2 cm hole in the bench and secured with a staple. The lower end of each tube featured two drainage holes, each 3 mm in diameter, which were connected to drainage pipes. To ensure darkness for the roots, a black plastic curtain was placed around the table, and a foam plastic plug was used to seal the hole, thereby blocking light and limiting airflow from beneath.

Two days before sowing, the tubes were connected to a watering drip-irrigation system to ensure adequate sand moisture for plant growth. Seeds were germinated on wet filter papers in petri dishes at room temperature (20°C) in a dark room for two days. Uniform sized pre-germinated seeds were selected, and a single seed was planted at a depth of 1 cm in the sand of each tube. A commercial plant nutrition solution (Wallco Växtnäring, Sweden) was delivered via an automatic watering system with a dosage of 2 ml per liter on a fixed schedule, ensuring a controlled water and nutrient regime throughout the experiment.

### Plant exposure to the volatiles from another cultivar

Six days after the initial sowing, individual plants in the growth stage of Zadoks decimal code 10-11 (Zadoks *et al*., 1974), were enclosed using the transparent twin-chamber cages, as shown in Fig. S1. These cages, placed on the table within the growth chamber, ensured the spatial separation and the containment of paired plants, while enabling the exposure of one cultivar to volatiles emitted by another cultivar. The system comprises a series of Perspex twin-chamber cages (10 x 10 x 80 cm), and each cage servs as a replicate. According to Ninkovic (2003) air was introduced into one chamber, designated as the inducing chamber (IC), through an opening (0.7 cm Ø), and then passed through an opening (0.7 cm Ø) in the partition wall into the adjacent responding chamber (RC). The air from the RC was directed into a vacuum tank before being vented outside the growing chamber. This design ensured that only aboveground volatiles were exchanged between the chambers, thereby excluding any influence of belowground compounds. The airflow through the system was 1.3 l min^-1^.

### Experimental treatments

We conducted two separate experiments: one with slow-growing cultivar Fairytale as the receiver plant; and the other with fast-growing cultivar Salome as the receiver. When Fairytale was used as receiver, the treatments were: Fairytale exposed to Fairytale (FeF), Fairytale exposed to Luhkas (FeL), and Fairytale exposed to Salome (FeS), with 16 replicates per treatment. In the Salome experiment, the treatments were: Salome exposed to Salome (SeS), Salome exposed to Luhkas (SeL), and Salome exposed to Fairytale (SeF), with 12 replicates per treatment. The treatments were randomly assigned and spatially arranged into 16 and 12 blocks, respectively, using a randomized block design to minimize environmental gradients and positional effects.

### Plant sampling and processing

On the 25^th^ day after planting, a time point at which volatile-mediated interactions between different cultivars typically produce the most pronounced differences in total dry weight (Ninkovic, 2003), plants were harvested for plant trait measurements and gene expression analyses. Plant samples were collected at early tillering growth stage, corresponding to decimal code 20–22 of the Zadoks scale (Zadoks *et al*., 1974). The plants were carefully removed by cutting the plastic sand tubes longitudinally and subsequently separated into two groups: roots and aboveground parts. Roots were gently washed with water to remove any adhering sand particles. The cleaned roots of each single plant were stored in a plastic bottle (100 ml) containing 10% alcohol in water solution, until size parameters were measured (max. 10 days). The detached leaves were carefully flattened using a book as a press. Leaves and roots were scanned using the image analysis system WinRHIZO Pro V 2007 (Regent Instruments, Québec, Canada). After scanning the leaves and roots, all plant samples (leaves, stems and roots) were dried separately in an oven at 70°C for 48 hours and then spent 24 h at room temperature before they were weighed.

### Morphological parameters

To evaluate whether receiver plants alter their growth strategy in response to volatiles emitted by cultivars with different growth strategies, we measured the total dry biomass of both emitter and receiver plants. Specifically, we examined the biomass of slow-growing Fairytale and fast-growing Salome receivers exposed to volatiles from each of the three cultivars. A total of 15 plant traits were recorded or calculated, with particular focus on parameters related to plant development and growth. These include biomass production parameters, such as dry biomass for leaves, stems, and roots, each measured separately after drying to constant weight. The calculated values for aboveground biomass (sum of leaf and stem biomass) and total biomass (sum of leaf, stem and root biomass) were also included.

Additional growth parameters included mass fractions of leaf (LMF), root (RMF), and shoot (SMF), which were determined by dividing the dry mass of leaves, roots, or shoots by the total plant dry mass, respectively. Shoot-to-root ratio (S/R) was calculated as the ratio of aboveground dry mass to root dry mass. Plant and stem heights were measured individually. Leaf surface area, representing the total surface area of all the leaves on a plant, was measured using the scanning and analysis software WinRhizo. Specific leaf area (SLA) was derived as the ratio of leaf surface area to leaf dry mass (m^2^ kg^−1^). For root traits, total length and average root diameter were quantified using root scanning and analysis software WinRhizo. While the dry biomass measurements served as the primary focus of this study, these supplementary morphological attributes provided additional insights into the overall plant growth and development in response to the neighbor volatiles.

### RNA high-throughput library preparation and bioinformatic analysis

At the end of the 25-day experiment, 1 cm long samples were collected from the tips of the youngest leaf of each receiver plant and immediately placed in Eppendorf tubes and kept in liquid nitrogen. From each treatment, three composite samples were generated by pooling leaf tips from seven biological replicates per sample. This resulted in nine pooled samples for RNA extraction and sequencing.

Total RNA was extracted using TRIzol reagent (Life Technologies) following the manufacturer’s instructions. mRNA for RNA sequencing was obtained by purification with the NEB mRNA isolation kit (New England Biolabs, Ipswich, MA, USA). Purified mRNA was used as input to prepare high-throughput RNA-seq libraries using the NEBNext Ultra II Directional RNA Library Prep Kit for Illumina (New England Biolabs). The resulting sequences were de-multiplexed, adapter trimmed, and quality controlled using TrimGalore (Krueger, 2015) (version 0.6.10) and FastQC (Andrews, 2010) (version 0.11.9). For gene expression analysis, RNA sequencing paired reads were aligned to the barley genome (Morex, version 3) using STAR (Dobin *et al*., 2013) (version 2.7.11b), with default parameters. HTSeq-count was used to count reads per gene with the parameters -mode union -stranded no -minequal 10 and -nonunique none. Count tables obtained were used in DESeq2 (Love *et al*., 2014) to infer significant expression with fit type set to parametric. All these tools were used through the Galaxy platform (Afgan *et al*., 2018).

### Plant volatile sampling and analysis

For the headspace analysis, twelve pre-germinated seeds were transplanted into pots (9 x 9 x 7 cm) containing planting soil (P-jord, Örebro, Hasselfors Garden). To prevent unwanted volatile interactions between the three cultivars, each pot was placed in a transparent plastic cage (10 x 10 x 80 cm), from which air was pumped out of the chamber (see description exposure system in the following). After 14 days of growth in the pots, reaching growth plant stage 12 of the Zadoks scale (Zadoks *et al*., 1974), plant VOCs were sampled and analysed.

To compare the volatile profiles from barley cultivars used as emitters, the headspace from potted Salome, Luhkas and Fairytale plants was sampled on adsorption tubes with a push-pull system for 24 h and analysed by gas chromatography-mass spectrometry (GC-MS). The upper part of each pot, containing 12 barley plants, was enclosed with a polyethylene terephthalate oven plastic bag (35 x 43 cm, Melita, Minden, Germany). Charcoal-filtered air was pushed into the oven bags with a flow of 600 mL min^-1^while pulling the air out of the bags over a Tenax TA sample tube (80 mg, 60/80 mesh size, GLScience, Eindhoven, Netherlands) with 400 mL min^-1^. Volatiles were released from the adsorbent tubes by thermal desorption with an Optics 3 Injector (GLScience, Eindhoven, Netherlands) at 250 °C. The thermal desorbed compounds were separated using an Agilent 7890N GC system equipped with a HP-1MS capillary column (30 × 0.25 mm id × 0.25 μm film thickness, 100% Dimethylpolysiloxane) coupled to Agilent 5975C mass spectrometer (Agilent Technologies, Inc., Santa Clara, CA, USA). Injection was employed using helium as carrier gas (Helium 6.0) with a flow of 1.3 mL min^-1^. The GC temperature program was as follows: Initial oven temperature of 30 °C was held for 2 min, increased at a rate of 5 C/min to 150 °C, followed by a rate of 10 C/min to the final temperature of 250 °C and held for 15 min. The GC inlet line temperature was 250 °C, and the ion source temperature was 180 °C. The quadrupole mass detector was operated in the electron impact (EI) mode at 70 eV, MS gain was set to 10. All data were obtained by collecting the full-scan mass spectra within the range of 40–500 m/z.

The volatile compounds from the chromatograms were identified and quantified with the “Automated Mass spectral Deconvolution and Identification System” (AMDIS, V. 2.71; National Institute of Standards and Technology NIST, Boulder, CO) following previously described methods (Gross *et al*., 2019). Identification criteria were applied as follows: match factor ≥ 80% and relative retention index deviation ≤ 5% + 0.01 from the reference value. The settings for deconvolution were component width, 32; adjacent peak subtraction, one; resolution, low; sensitivity, medium; shape requirements, low; level, very strong; maximum penalty, 25; and “no RI in library” 20. Components with a signal to noise ratio < 300 were excluded from the analysis.

### Statistical analysis

Statistical analyses and plotting of morphological attributes, such as total biomass production, were conducted using R (version 4.3.0) in the RStudio environment (version 2024.09.1+394, ’Cranberry Hibiscus’). A two-way ANOVA was performed to identify significant differences among treatment groups at a 95% confidence level. The model included volatile treatment (i.e., identity of the neighboring plant cultivar) and block (i.e., experimental position within the growth chamber) as fixed factors. Interactions between treatment and block were initially tested but found to be non-significant and were excluded from the final model. Block was included to control for spatial variation rather than to test for its direct effect. When a significant main effect of treatment was detected (*p* ≤ 0.05), Tukey’s HSD post-hoc tests were used to assess pairwise differences among treatments.

Differential expression analysis of the epigenetic measurements was conducted using DESeq2 (version 1.44.0) in the R statistical environment (version 4.4.0). Significantly differentially expressed genes were identified based on appropriate statistical cutoffs, such as a false discovery rate (FDR) threshold of <0.05 and log2 fold change (log2FC) values < -1 or > 1.

Random Forest (RF) approach was used to compare the composition of plant VOCs released from the barley cultivars. Ntree=10.000 bootstrap trees were drawn with myrt=11 random variables at each node. A confusion matrix of the RF model and the corresponding classification errors were used to investigate the assignment accuracy of individual samples to the different cultivars. To identify the most important compounds responsible for the resolution of samples, the average mean decrease in accuracy (MDA) was calculated with the *“importance”* function from the *randomForest* package (Liaw & Wiener, 2002).

The amount (peak area in chromatogram) of the ten most important VOCs that lead to the separation of the volatile profiles from the different barley cultivars was compared using Kruskal-Wallis tests. Followed by pairwise comparison of cultivars with Dunn’s test using the *“DunnTest”* function from the *FSA* package (Ogle *et al*., 2015), *p*-values of multiple comparisons were adjusted with the Bonferroni method.

## Results

### Constitutive VOCs affect biomass production of divergent barley cultivars

To test whether constitutive VOCs modulate biomass production in neighboring plants, we conducted two independent exposure experiments using barley cultivars with contrasting growth rates: Fairytale (slow), Luhkas (intermediate), and Salome (fast). In each experiment, either Fairytale or Salome acted as the receiver. When Fairytale was the receiver, it was exposed to VOCs emitted by Fairytale (FeF), Luhkas (FeL), or Salome (FeS). Emitter biomass differed significantly among cultivars in this experiment (Fig. 2a), with Fairytale producing less biomass than Luhkas (*p* = 0.006) and Salome (*p* < 0.001), and Luhkas producing less than Salome (*p* = 0.014). Receiver biomass (Fig. 2b) was also influenced by emitter identity (ANOVA: *F* (2, 30) = 3.97, *p* = 0.029). Fairytale plants exposed to constitutive VOCs from Salome had significantly higher biomass than those exposed to VOCs from Fairytale emitters (*p* = 0.025), while exposure to Luhkas did not differ significantly from the other treatments (*p* > 0.05).

**Fig. 2.**
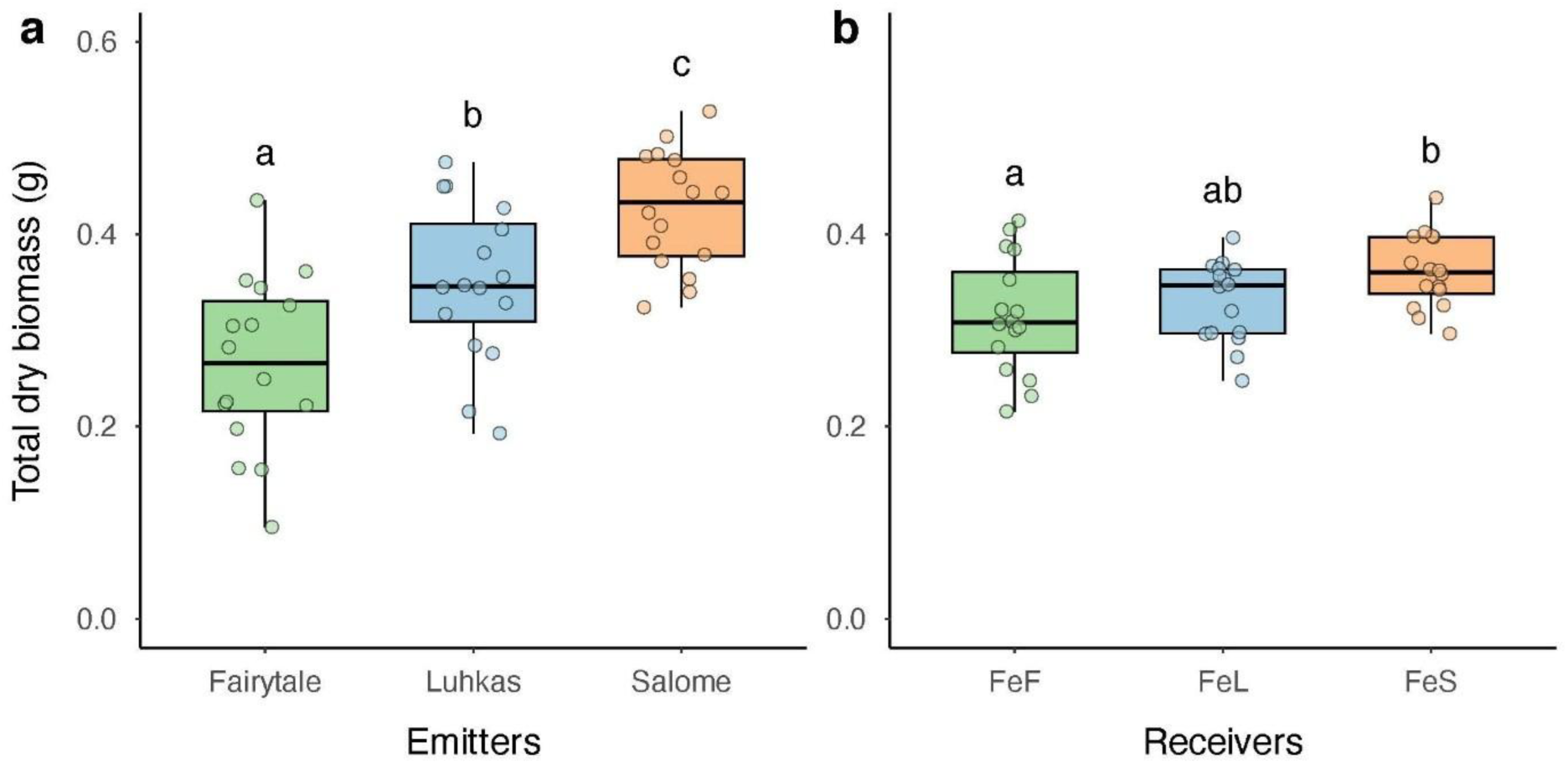
Biomass responses in Fairytale receiver plants and emitters. Total dry biomass of (a) emitter plants and (b) Fairytale receiver plants after exposure to volatiles from different cultivars: FeF = Fairytale exposed to Fairytale, FeL = Fairytale exposed to Luhkas, FeS = Fairytale exposed to Salome. Box plots show the interquartile range (IQR), median (horizontal line), and whiskers extending to 1.5× IQR. Each point represents one plant (n = 16). Different letters indicate statistically significant differences (p ≤ 0.05, two-way ANOVA with Tukey’s HSD post hoc test).

In the second exposure experiment, the fast-growing cultivar Salome served as the receiver and was exposed to constitutive VOCs emitted by Salome (SeS), Luhkas (SeL), or Fairytale (SeF). As in the first experiment, emitter biomass differed significantly among cultivars (Fig. 3a). Fairytale emitters produced less biomass than Luhkas (*p* = 0.032) and Salome (*p* < 0.001), while Luhkas and Salome did not differ significantly (*p* > 0.05). As shown in Fig. 3b, the biomass of Salome as the receiver was also significantly influenced by emitter identity in this experiment (ANOVA: *F* (2, 22) = 9.74, *p* < 0.001). Salome plants exposed to constitutive VOCs from Fairytale had lower biomass than those exposed to Salome (*p* < 0.001). Exposure to Luhkas also reduced biomass relative to Salome (*p* = 0.046). No significant difference was found between exposure to Fairytale or Luhkas (*p* > 0.05). Together, these results demonstrate that constitutive VOCs can modulate biomass accumulation in neighboring plants. VOCs from fast-growing cultivars (Salome) enhance receiver growth, while VOCs from slow-growing cultivars (Fairytale) suppress it.

**Fig. 3.**
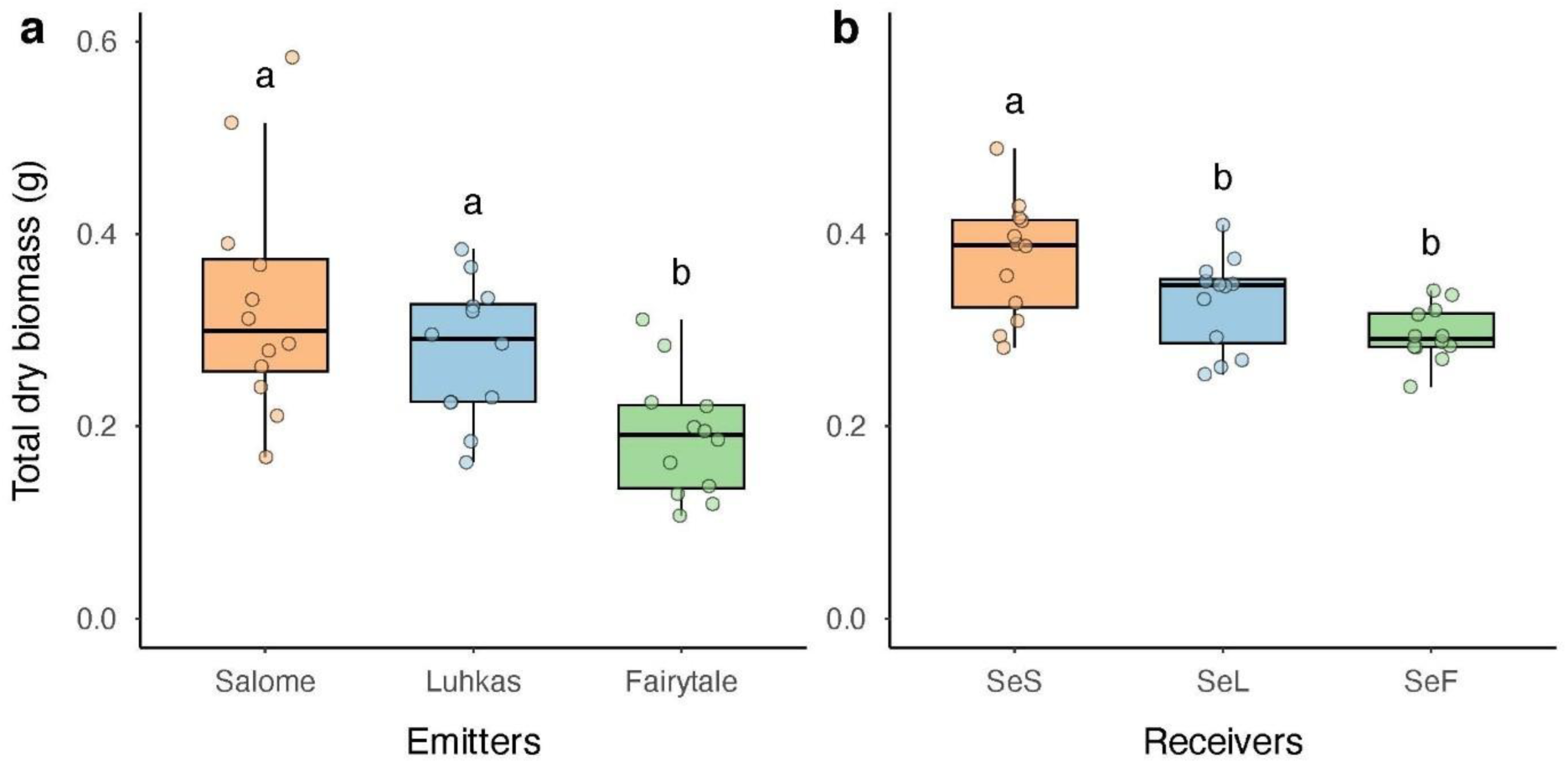
Biomass responses in Salome receiver plants and emitters. Total dry biomass of (a) emitter plants and (b) Salome receiver plants after exposure to volatiles from different cultivars: SeS = Salome exposed to Salome, SeL = Salome exposed to Luhkas, SeF = Salome exposed to Fairytale. Box plots show the interquartile range (IQR), median (horizontal line), and whiskers extending to 1.5× IQR. Each point represents one plant (n = 12). Different letters indicate statistically significant differences (p ≤ 0.05, two-way ANOVA with Tukey’s HSD post hoc test).

### Additional morphological traits show limited response to VOC exposure

Beyond total biomass, we assessed a range of morphological and allocation traits to determine whether VOC exposure affected other aspects of plant development. In the experiment where Fairytale was the receiver, no significant differences were observed among treatments for leaf, shoot, or root mass fractions (Table S1). The shoot-to-root ratio was significantly higher in Salome emitters compared to Fairytale (*p* = 0.034) and Luhkas (*p* = 0.036), while receivers showed no differences. Leaf surface area increased significantly across emitter cultivars from Fairytale to Luhkas to Salome, with all pairwise comparisons being significantly different (F– L = 0.008, F–S < 0.001, L–S < 0.001). In contrast, no differences were observed among receiver plants. Specific leaf area (SLA), plant height, stem heights and average root diameter did not vary significantly across treatments. Root length increased significantly across emitter cultivars (F–L = 0.017, F–S < 0.001, L–S = 0.042), while no significant differences were detected in receivers. In summary, emitter identity had no discernible effect on any of the additional traits measured in Fairytale receivers.

In the experiment where Salome was the receiver, leaf and shoot mass fractions were significantly lower in Luhkas emitters compared to Salome, with Fairytale showing intermediate values (Table S2). The difference between Luhkas and Salome was significant for both leaf mass fraction (*p* = 0.034) and shoot mass fraction (*p* = 0.002). Receiver plants showed no significant differences in leaf or shoot mass fractions. Root mass fraction and shoot-to-root ratio exhibited no significant treatment effects. Leaf surface area increased significantly among emitter cultivars, from Fairytale to Luhkas to Salome (F–L = 0.04, F–S < 0.001, L–S = 0.015). Among receivers, SeF had a lower leaf surface area than SeS (*p* = 0.004), with SeL being intermediate. SLA was largely consistent across treatments, though a slight but significant difference was detected between Luhkas and Salome emitters (*p* = 0.045). In terms of plant height, SeS was significantly taller than both SeL (*p* = 0.003) and SeF (*p* = 0.023), with no difference between SeF and SeL. For stem height, SeL was significantly shorter than both SeS (*p* < 0.001) and SeF (*p* = 0.047), whereas SeS and SeF did not differ significantly. Root length in emitter plants was significantly lower in Fairytale compared to Salome (*p* = 0.002) and Luhkas (*p* = 0.011). Among receivers, SeS had significantly greater root length than both SeF (*p* = 0.003) and SeL (*p* = 0.015). No significant treatment differences were observed in average root diameter. Overall, emitter identity altered only a subset of traits in Salome receivers, specifically leaf surface area, plant height, stem height, and root length, while all other measured parameters showed no significant or consistent treatment-related effects.

### Transcriptional response to constitutive VOC exposure mimics biomass production

The level of physiological changes observed in our experimental setup prompted us to explore the genome-wide transcriptional reprogramming taking place in receiver plants exposed to constitutive VOCs. To that aim we prepared and sequenced high-throughput mRNA libraries from the two exposure-experiments with a focus on receiver plants. Our differential expression analysis (adjusted *p*-value < 0.05) revealed significant differences in gene expression between treatments (Fig. 4a), as expected from a PCA of the RNA libraries (Fig. S2a,d). Fairytale exposed to Salome (FeS) exhibited a higher number of down-regulated genes (24 up-regulated and 69 down-regulated genes compared to its control Fairytale exposed to Fairytale, FeF). Similarly, when FeS was compared to FeL we observed 45 up-regulated and 61 down-regulated genes (Fig. S2b), indicating that FeS receiver plants show the higher transcriptional changes from all the Fairytale receiver combinations. As expected, FeL compared to FeF showed minimal transcriptional changes (Fig. S2c). Interestingly, Salome exposed to Fairytale (SeF) exhibited the opposite transcriptional response, with a higher number of up-regulated genes (2,172 up-regulated genes and 248 down-regulated genes compared to Salome exposed to Salome, SeS). This result was consistent when SeF receivers were compared to SeL, showing a similar tendency to gene up-regulation (2,422 up-regulated genes and 303 down-regulated genes) (Fig. S2e). Similar to the result observed in Fairytale used as receiver, the comparison between SeS and SeL revealed no significant differences at the transcriptional level (Fig. S2f).

**Fig. 4.**
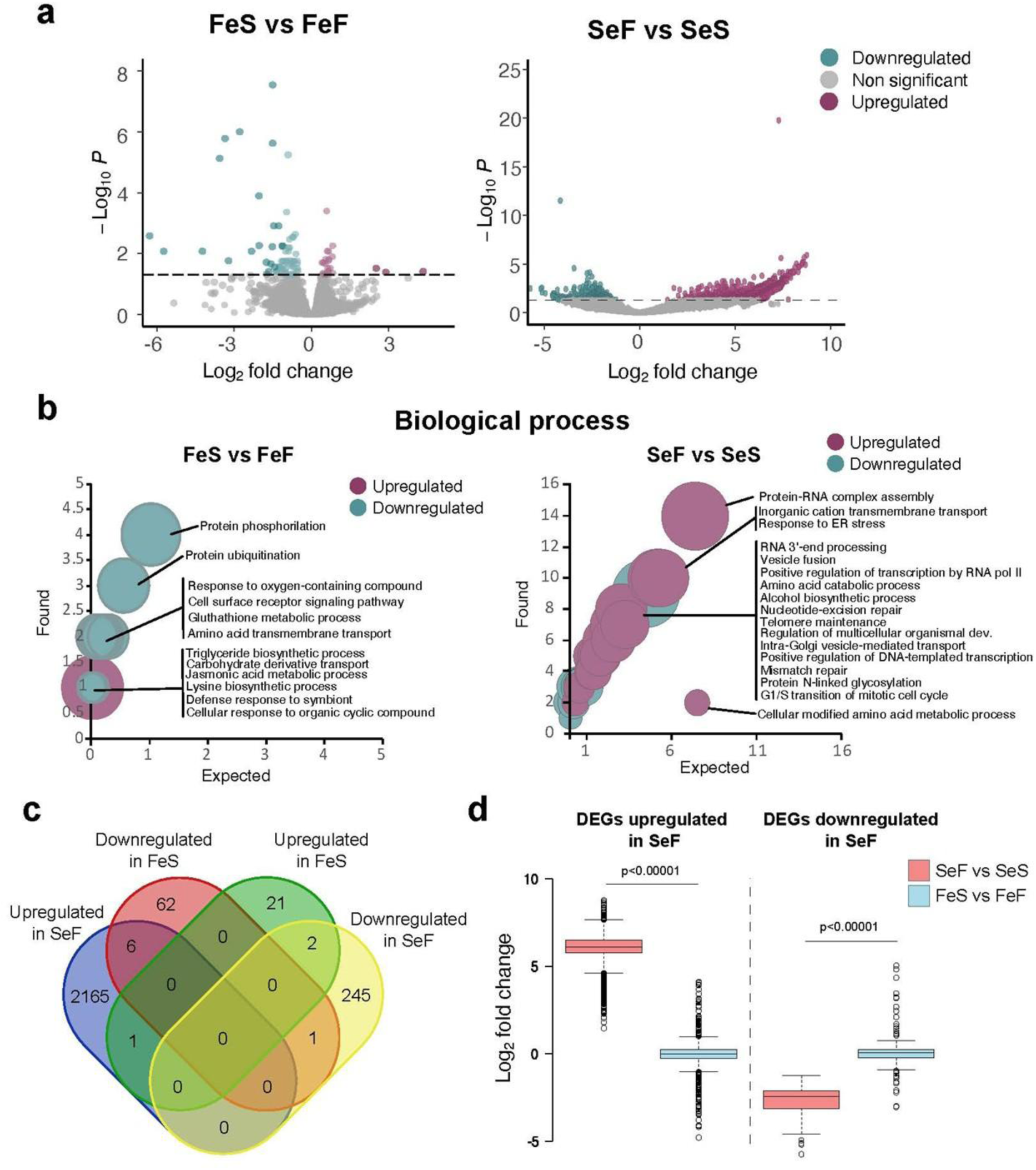
Gene expression responses to VOC exposure. (a) Volcano plots showing the up- and down-regulated genes in both experiments. Left panel: FeS compared to FeF. Right panel: SeF compared to SeS. (b) Gene ontology functional enrichment plots, showing biological processes categorization in the comparisons between FeS and FeF (left panel), and SeF and SeS (right panel). (c) Venn diagram showing the overlap between genes up- and down-regulated in the SeF and FeS. (d) Expression level of DEGs up- and down-regulated in SeF, in both the comparisons SeF vs SeS and FeS vs FeF. p-value was calculated through an unpaired, unequal variance t-test.

Gene ontology functional enrichment analyses (Fig. 4b) were conducted to provide a broad overview of the functional roles of the up- and down-regulated genes in the plants. In Fairytale exposed to Salome compared to Fairytale exposed to Fairytale (FeS vs. FeF), the primary functional categories (according to the biological process categorization) associated with the down-regulated genes were involved in protein homeostasis (protein phosphorylation, ubiquitination, amino acid transmembrane transport, triglyceride biosynthetic process, etc.) and the response to stress (response to oxygen-containing compound, defense response to symbiont, cellular response to organic cyclic compound, etc.). In comparison with Salome as the receiving cultivar, genes differentially expressed were associated with processes involved in RNA homeostasis (protein-RNA complex assembly, RNA 3’-end processing, regulation of transcription, etc.), cellular transport (inorganic cation transmembrane transport, vesicle fusion, intra-golgi vesicle-mediated transport, etc.), DNA replication (telomere maintenance, mismatch repair, G1/S transition of mitotic cell cycle), and protein homeostasis (amino acid catabolic process).

To further compare the transcriptional responses in the two receiver cultivars, we explored the overlap between up- and down-regulated genes on each comparison (Fig. 4c). This showed that there was a small overlap of differentially expressed genes (DEGs) up-regulated in SeF that were also down-regulated in the opposite situation FeS (6 genes, 8.7% of the total down- regulated DEGs in FeS). Similarly, a number of down-regulated genes in SeF were up-regulated in FeS (2 genes, 8.3% of the total up-regulated DEGs in FeS). Next, we explored the changes in total expression for SeF up- and down-regulated genes (Fig. 4d). Interestingly, we found that up-regulated DEGs in SeF have significantly lower expression levels in FeS, while down- regulated DEGs in FeS, showed significantly higher expression levels then in SeF (Fig. 4d). In summary, our transcriptomic analysis indicated that SeF and FeS experienced antagonistic transcriptional changes and a significant increase in the expression of genes associated with categories that explain their physiological changes after exposure to specific emitters.

### Constitutive VOC profiles differ between barley cultivars with divergent growth strategies

VOC composition varies both across plant species (Vivaldo *et al*., 2017) and among cultivars (Dahlin *et al*., 2018). To examine the differences among the three barley cultivars in this study, we analysed the headspace composition of 115 annotated volatile compounds using gas chromatography-mass spectrometry (GC-MS). A random forest model achieved 93.1% accuracy in classifying volatile profiles from undamaged plants. Constitutive VOC profiles were clearly separable among Salome, Fairytale and Luhkas (Fig. 5a), with Salome and Luhkas clustering more closely together than either did with Fairytale. All Luhkas samples were assigned correctly with no confusion with other cultivars (Fig. 5b). Salome VOC profiles were classified with 90% accuracy, a single sample was misclassified as Luhkas (Fig. 5b). Eight out of 9 volatile samples from Fairytale were correctly classified as Fairytale, while one was wrongly assigned as Luhkas (Fig. 5b). Notably, VOC profiles from Salome and Fairytale were consistently distinguishable, showing the greatest separation in both clustering and classification (Fig. 5a,b).

**Fig. 5.**
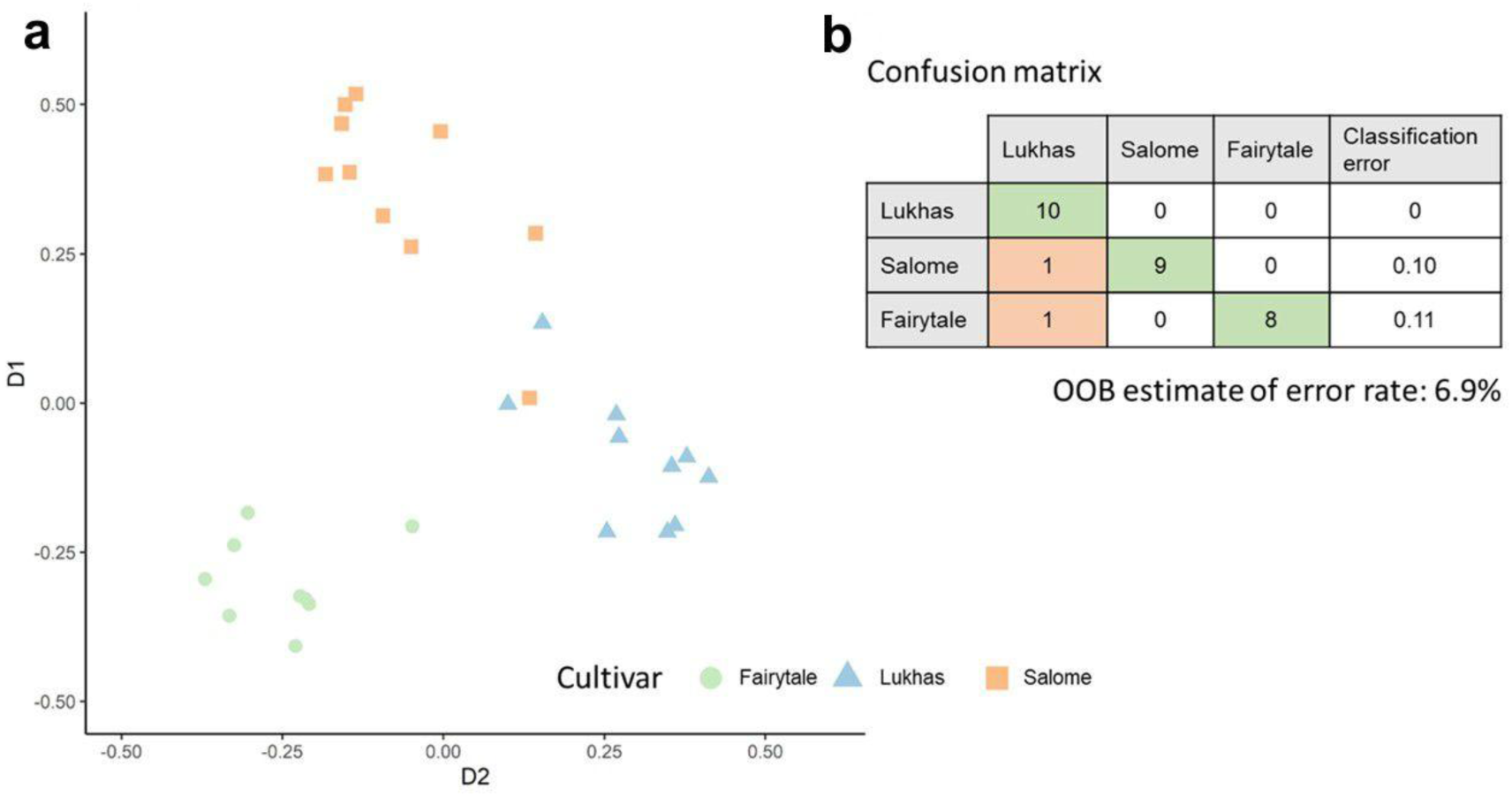
Classification of cultivar-specific VOC profiles. (a) Multi-dimensional scaling (MDS) plot of random forest model using the proportion of volatile organic compounds released from barley cultivars Fairytale (green circles, n = 9), Luhkas (blue triangles, n = 10), and Salome (red squares, n = 10). (B) Corresponding confusion matrix of sample classification of the RF model. The matrix shows the correct (green) and false (red) assignment of cultivar VOC profiles and the classification error for each cultivar.

The ten VOCs, which had the greatest influence on the separation of the cultivars (MDA value), were emitted in significantly different quantities from the three cultivars (Kruskal-Wallis, df = 2, RI 1160: χ2 = 18.26, *p* < 0.001; benzyl nitrile: χ2 = 18.99, *p* < 0.001; 1-octen-3-ol: χ2 = 24.06, *p* < 0.001; benzothiazole: χ2 = 6.36, *p* = 0.042; tetradecane: χ2 = 8.12, *p* = 0.017; octanal: χ2 = 15.52, *p* < 0.001; dodecane: χ2 = 8.43, *p* = 0.015; linalool: χ2 = 13.81, *p* = 0.001; RI 955: χ2 = 13.45, *p* = 0.001; nonanal: χ2 = 14.62, *p* < 0.001).

Benzyl nitrile and the unidentified compound RI 1160 were significantly more released from Fairytale compared to Luhkas and Salome (Fig. 6a). The release of 1-octen-3-ol was characteristic for Salome plants (Fig. 6a). Tetradecane, octanal and linalool were released in different amounts from Fairytale compared to Salome, while there was no significant difference from both cultivars compared to Luhkas (Fig. 6b). In chromatograms of samples from Fairytale and Luhkas significantly greater peak areas (amounts) of nonanal were detected compared to samples from Salome (Fig. 6b). In conclusion, each cultivar produced a distinct constitutive VOC profile, with key compounds varying in both identity and abundance. These differences likely underpin the cultivar-specific growth and transcriptional responses observed in receiver plants.

**Fig. 6.**
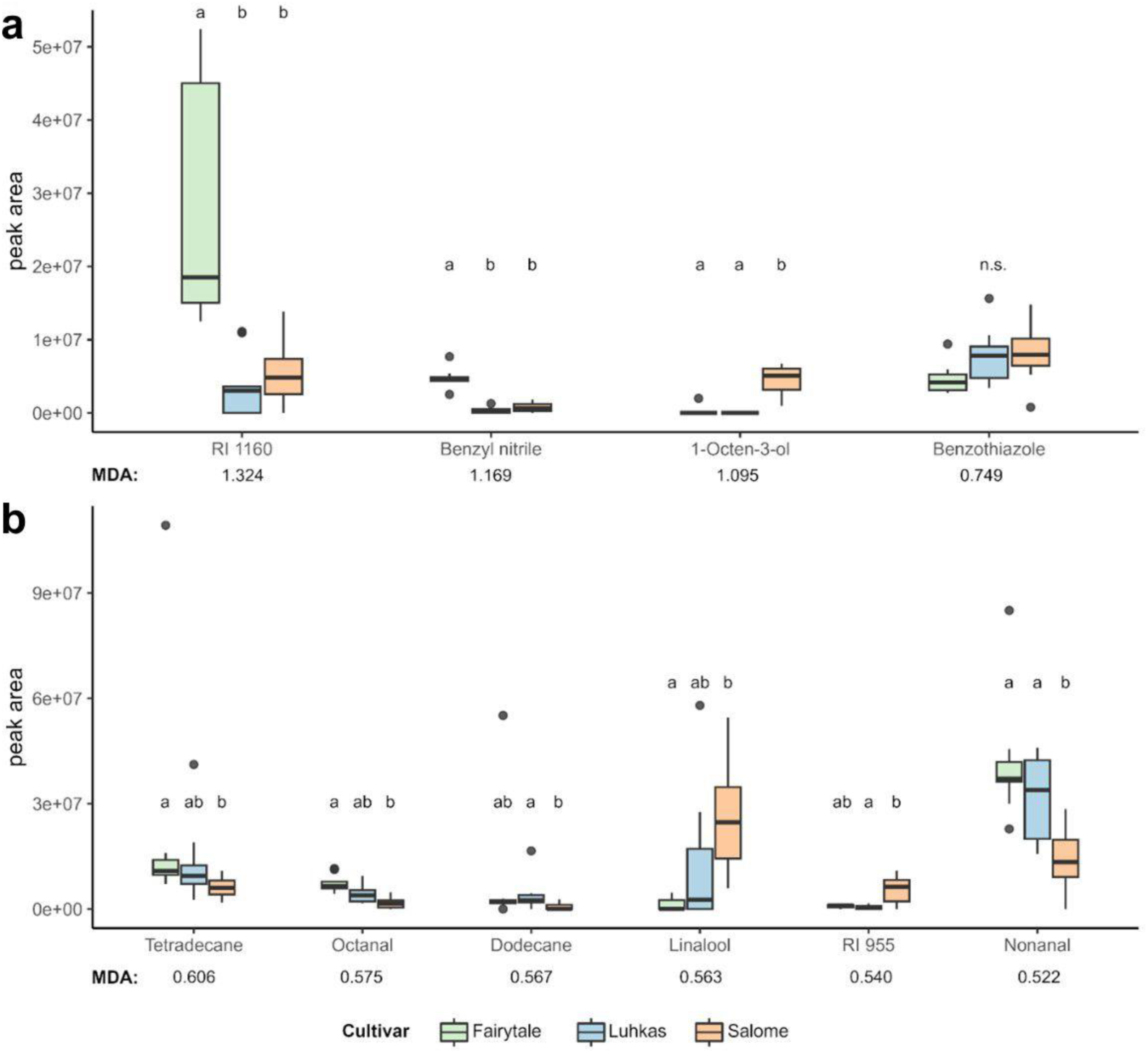
Most influential VOCs in cultivar classification. Peak area (amount) of VOCs released from three different barley cultivars: Fairytale (green, n = 9), Luhkas (blue, n = 10), and Salome (red, n = 10), which have the greatest influence on the separation of cultivars by the random forest model. Impact on the separation is evaluated by the mean decrease in accuracy (MDA). Boxes correspond to the 25^th^ and 75^th^ percentiles, medians are shown as lines, and whiskers extend to 1.5 times of the interquartile ranges. Dots represent outliers. Letters indicate significant differences (*p* < 0.05) of pairwise comparisons with Dunns’ Test with Bonferroni correction as post-hoc test after sig. Kruskal-Wallis rank sum test.

## Discussion

Undamaged plants emit distinct VOC profiles reflecting their inherent growth strategies (Ninkovic *et al*., 2016; Dahlin *et al*., 2018). Our findings demonstrate that neighboring plants can detect these constitutive VOCs and adjust their growth-defense allocation accordingly, aligning with the ecological principle that plants need to balance resources between growth and defense (Herms & Mattson, 1992; Züst & Agrawal, 2017). In our experiments, the slow-growing cultivar Fairytale increased biomass when exposed to the fast-growing Salome (FeS), whereas Salome showed reduced biomass when exposed to Fairytale (SeF). The intermediate- growing Luhkas consistently showed moderate responses. This pattern supports the context-dependence of plant responses to neighbor VOCs, a concept reflected in studies reporting both growth stimulation and inhibition following exposure to herbivore-induced plant volatiles (HIPVs) (Cofer *et al*., 2018; Freundlich *et al*., 2021). The extent and direction of the observed biomass changes were closely associated with the differences in growth strategies between VOC emitters and receivers, underscoring plants’ ability to interpret subtle chemical cues and adjust allocation accordingly. Gene expression analyses reinforced this trend, with more differentially expressed genes (DEGs) detected in interactions between Fairytale and Salome compared to those involving Luhkas. These DEGs were largely linked to cell growth and defence activation, mirroring previous observations that up-regulation of resistance genes can compromise growth (Cipollini & Heil, 2010).

These findings reflect adaptive resource allocation strategies consistent with growth-defense trade-offs (Herms & Mattson, 1992; Züst & Agrawal, 2017). Fast-growing plants typically invest in rapid biomass accumulation over defense, while slow-growing ones allocate more to defense (Herms & Mattson, 1992). Therefore, exposure to a slow-growing neighbor may convey lower competitive pressure and prompt defense investment, while exposure to a fast-growing neighbor may elicit a growth-oriented response, potentially at the cost of induced resistance. Similar effects have been reported for induced VOCs released by damaged plants, such as in *Artemisia tridentata,* which increases resistance but reduces growth in response to neighbor volatiles (Karban, 2017), or in specific HIPVs like (Z)-3-hexenyl acetate that mediate growth-defense trade-offs (Engelberth & Engelberth, 2019; Freundlich *et al*., 2021). However, unlike these stress-induced signals, our findings demonstrate that such physiological shifts can also be triggered by constitutive VOCs emitted by undamaged plants.

The three cultivars in our study emitted distinct VOC profiles, with the greatest difference between Salome and Fairytale. This chemical dissimilarity likely drives the adaptive responses in receiver plants, contributing to shifts in growth-defense trade-offs. Benzyl nitrile and a chemically unidentified compound (retention index 1160) were characteristic of Fairytale’s VOC profile. The emission of benzyl nitrile, a well-known HIPV with insect-repellent properties (Dong *et al*., 2011; Twidle *et al*., 2022; Qian *et al*., 2023), may reflect a defense-oriented strategy in Fairytale. Although its role in interactions among undamaged plants has not yet been investigated, it may function as a cue that alters growth-defense allocation in neighboring plants. Tetradecane and octanal also emerged as potentially significant compounds, as their emission levels varied sharply across cultivars. This supports the idea that specific VOC concentrations are required to pass thresholds and induce a physiological response in neighboring plants, a concept consistent with the functioning of chemoreceptors in plants (Boller, 1995; Wicher, 2018). Given that defense pathways can be induced in undamaged receivers, and considering the reciprocal trade-off between growth and defense, we speculate that such induced responses may significantly impact competitive dynamics. Since competition is a major stressor for plants (Goldberg & Barton, 1992), VOC-mediated growth-defense adjustments may represent an adaptive strategy that enable plants to anticipate and prepare for future competitive challenges, analogous to how HIPVs prime defenses against herbivores. In this context, constitutive VOCs may act as priming signals, orienting receiver plants toward growth or defense depending on the perceived competitive strength of their neighbors.

Gene expression analyses showed that plants actively adjust their transcriptional programs in response to constitutive VOCs emitted by undamaged neighbors, with distinct patterns emerging between cultivars. When Fairytale was exposed to Salome (FeS), most DEGs were down-regulated, particularly those related to protein maintenance, stress responses, and phosphorylation, suggesting reduced investment in defense. In contrast, Salome exposed to Fairytale (SeF) showed broad up-regulation of DEGs associated with RNA processing, DNA replication, and protein metabolism, potentially indicating increased readiness for stress or development. Notably, in SeF conditions, WRKY transcription factors associated with enhanced resistance to aphids (Wani *et al*., 2021) were up-regulated, aligning with previous findings that VOC exposure can reduce aphid performance (Dahlin *et al*., 2018; Kheam *et al*., 2023). Interestingly, some DEGs shower opposite regulations in FeS and SeF conditions. For example, a universal stress protein (Chi *et al*., 2019) and a receptor-like serine/threonine kinase (Brueggeman *et al*., 2002, 2006; Nirmala *et al*., 2006), key players in biotic stress response, were up-regulated in SeF but down-regulated in FeS. Other genes such as a WAK family protein (Tripathi *et al*., 2021) and an AB hydrolase-1 domain protein (Zakhrabekova *et al*., 2023) may influence broader development and stress responses. In addition, a RING-type E3 ubiquitin ligase (You *et al*., 2016) may contribute to disease resistance and the coordination of growth-defense trade-offs. Conversely, two genes up-regulated in FeS but down-regulated in SeF include a predicted protein and a V-ATPase involved in turgor regulation and growth (Li *et al*., 2022). These contrasting patterns underscore cultivar-specific responses and support the hypothesis that VOC perception shapes physiological priorities in growth and defense.

Perception of VOCs is essential for optimizing plant fitness through rapid and fine-tuned physiological adjustments (Ninkovic *et al*., 2021). This supports the hypothesis that growth-defense trade-offs are not static, but context-dependent and dynamically regulated in response to neighbor cues. Plants may use VOCs to assess the competitive ability of their neighbors and adapt their own strategies within the plant community. Exposure to a fast-growing, high-competition neighbor prompts investment in growth at the expense of defense, whereas exposure to a slow-growing, low-competition neighbor may reduce growth and promote allocation toward induced defenses. Such strategic adjustments resemble those seen in response to herbivory (Engelberth *et al*., 2004), pathogen presence (Hammerbacher *et al*., 2019), or environmental stress (Caparrotta *et al*., 2018; Markovic *et al*., 2019), all of which can modulate VOC production and prepare neighboring plants for future challenges. Our findings demonstrate that even constitutive VOC emissions from undamaged plants can serve as potent ecological signals that influence the growth strategies of neighboring plants.

## Acknowledgements

The authors gratefully acknowledge Maria Hedsén, Ahmed Ez, and Emmanuel Hiltcher for their technical assistance and support during the course of this research. We appreciate the financial support by the European Union’s Horizon 2020 Research and Innovation Programme through the EcoStack project (Grant Agreement no. 773554).

## Author contributions

VN planned and designed the research, AA, MR, JG and DM performed and conducted experiments, AA, ID, JG, GM and TG analyzed data, AA, MR, JG, GM, DM and VN wrote the manuscript, AA and MR contributed equally.

