## Supplementary material for "Volatile compounds released by undamaged plants influence the adaptive growth strategies of neighboring plants": Fig S1

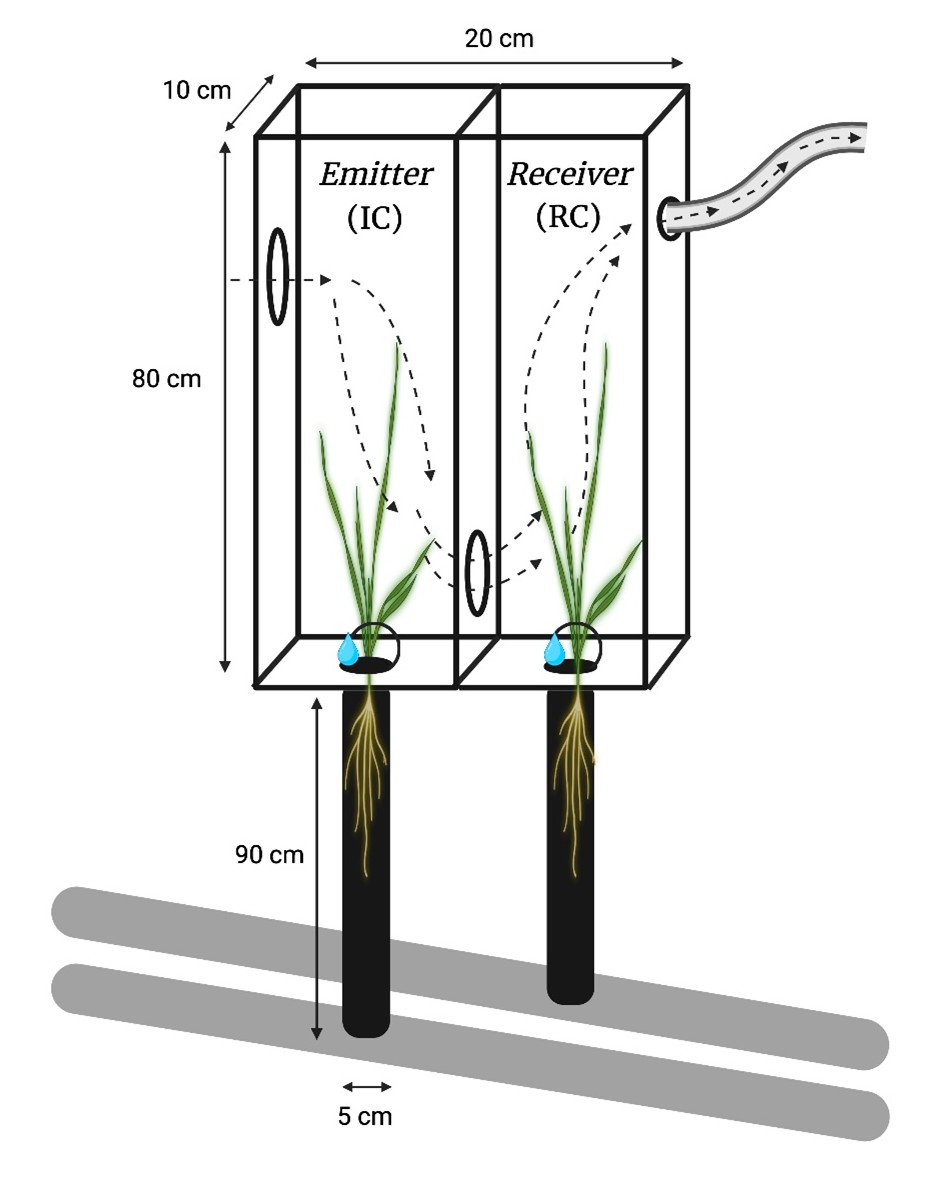

**Fig. S1** Twin-chamber volatile-exposure system. An emitter plant is grown in the inducing chamber (IC), while a receiver plant is placed in the responding chamber (RC). Air is drawn through a vent in the upper right corner of the responding chamber, creating airflow from the opening in the inducing chamber. Each chamber is equipped with an individual drip-irrigation system. Plastic tubes filled with sand are placed below the chambers to support root growth, with their lower ends connected to a drainage pipes. Dashed arrows indicate airflow and the movement of volatile organic compounds through the chambers. (Created with Biorender.com).
