## Supplementary material for "Volatile compounds released by undamaged plants influence the adaptive growth strategies of neighboring plants": Fig S2

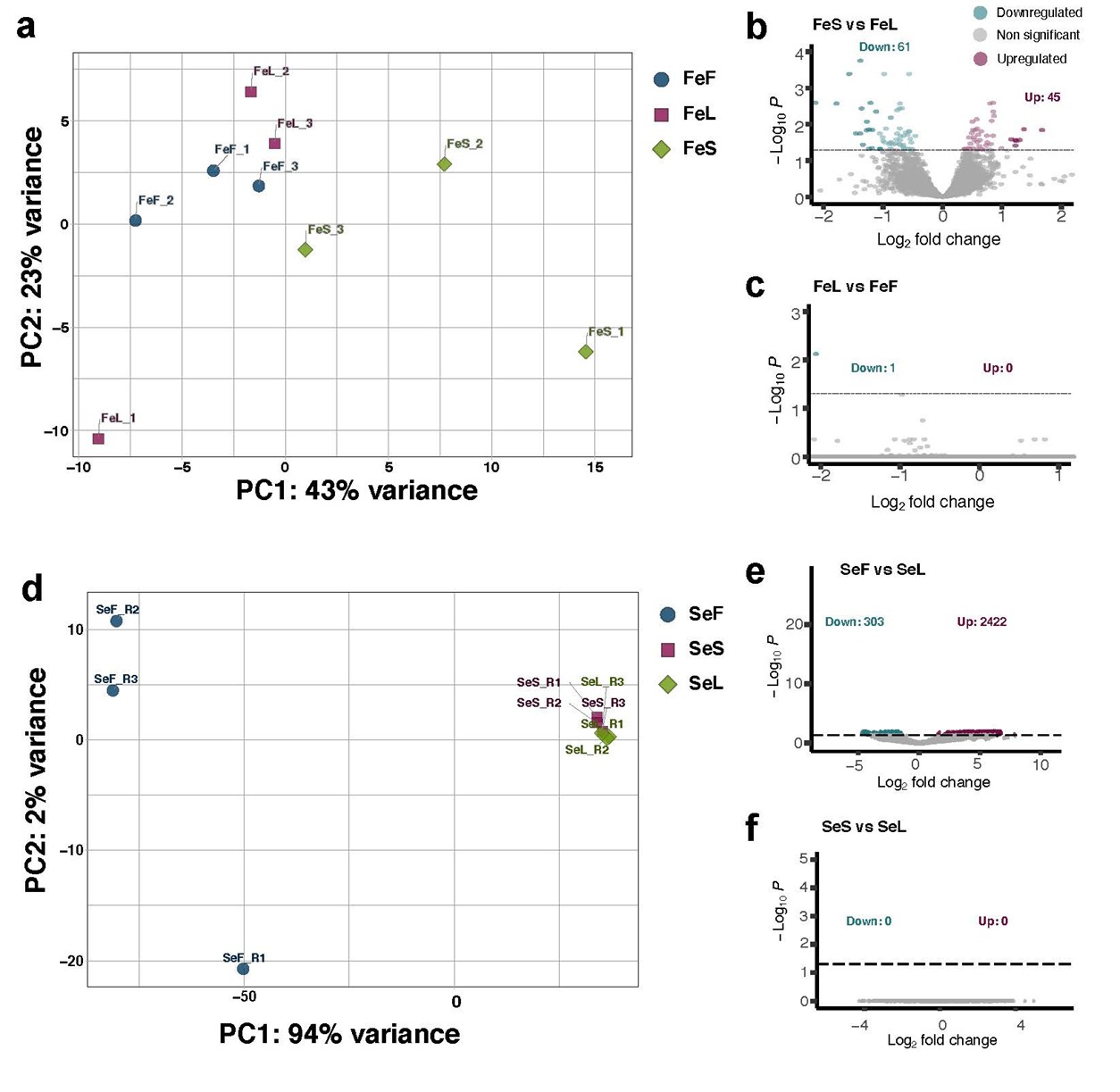

**Fig. S2** **a)** Principal component analysis of RNA libraries generated from Fairytale receiver combinations. Numbers represent bioreplicates analyzed for each combination. **b**-**c)** Volcano plots showing the up- and down-regulated genes in FeS compared to FeL (**b**), and FeL compared to FeF (**c**). **d)** Principal component analysis for the RNA libraries produced from Salome receiver combinations. Numbers indicate bioreplicates analyzed for each combination. **e**-**f)** Volcano plots showing the up- and down-regulated genes in SeF compared to SeL (**e**), and SeS compared to SeL (**f**).
