## Supplementary material for "Volatile compounds released by undamaged plants influence the adaptive growth strategies of neighboring plants": Table S2

| **Table S2.** Overview of measured plant traits (mean ± SE) in Salome, exposed to volatiles from different emitter cultivars (Salome, Luhkas, or Fairytale): Salome exposed to Salome (SeS), Salome exposed to Luhkas (SeL), Salome exposed to Fairytale (SeF). Columns represent treatment means with superscript letters (a, b, c) indicating statistically significant differences among groups (*p* < 0.05, Tukey’s HSD). Statistical comparisons are based on two-way ANOVA with treatment and block as fixed factors. | | | | | | | |
| --- | --- | --- | --- | --- | --- | --- | --- |
|  |  | Emitters |  |  |  | Receivers |  |
| Plant trait | Salome | Luhkas | Fairytale |  | SeS | SeL | SeF |
| Leaf mass fraction (g/g) | 0.51^a^  ± 0.012 | 0.48^b^  ± 0.009 | 0.49^ab^  ± 0.007 |  | 0.52^a^  ± 0.014 | 0.51^a^  ± 0.013 | 0.52^a^  ± 0.006 |
| Shoot mass fraction (g/g) | 0.29^a^  ± 0.005 | 0.34^b^  ± 0.007 | 0.31^ab^  ± 0.009 |  | 0.3^a^  ± 0.009 | 0.29^a^  ± 0.009 | 0.3^a^  ± 0.009 |
| Root mass fraction (g/g) | 0.2^a^  ± 0.01 | 0.19^a^  ± 0.007 | 0.19^a^  ± 0.008 |  | 0.18^a^  ± 0.012 | 0.2^a^  ± 0.007 | 0.18^a^  ± 0.007 |
| Shoot-to-root ratio | 4.12^a^  ± 0.22 | 4.48^a^  ± 0.21 | 4.22^a^  ± 0.2 |  | 4.75^a^  ± 0.41 | 4.13^a^  ± 0.2 | 4.61^a^  ± 0.32 |
| Leaf surface area (cm^2^) | 57.8^a^  ± 3.03 | 44..5^b^  ± 3.25 | 32.2^c^  ± 2.23 |  | 70.9^a^  ± 2.8 | 64.3^ab^  ± 2.95 | 58.7^b^  ± 1.08 |
| Specific leaf area (cm^2^g^-1^) | 371^a^  ± 22.7 | 339^b^  ± 14.7 | 366^ab^  ± 16.9 |  | 370^a^  ± 13.6 | 385^a^  ± 11.7 | 388^a^  ± 10.2 |
| Plant height (cm) | 47.2^a^  ± 0.42 | 48.1^a^  ± 0.67 | 46.5^a^  ± 0.55 |  | 50.5^a^  ± 1.02 | 46.4^b^  ± 0.99 | 47.4^b^  ± 1.02 |
| Stem height (cm) | 13.7^a^  ± 0.67 | 15.2^a^  ± 0.55 | 15.1^a^  ± 0.38 |  | 14.5^a^  ± 0.42 | 12.1^b^  ± 0.42 | 13.5^b^  ± 0.42 |
| Root length (cm) | 1345^a^  ± 75.8 | 1288^a^  ± 55.7 | 1032^b^  ± 69.9 |  | 1719^a^  ± 81.3 | 1490^b^  ± 62.4 | 1442^b^  ± 67.2 |
| Average root Ø (cm) | 0.33^a^  ± 0.012 | 0.31^a^  ± 0.014 | 0.31^a^  ± 0.016 |  | 0.31^a^  ± 0.009 | 0.33^a^  ± 0.007 | 0.32^a^  ± 0.009 |
